# Clinical characteristics and primary management of patients diagnosed with prostate cancer between 2015 and 2019 at the Uganda Cancer Institute

**DOI:** 10.1101/2020.07.09.194936

**Authors:** Paul Katongole, Obondo J. Sande, Mulumba Yusuf, Moses Joloba, Steven J Reynolds, Nixon Niyonzima

**Affiliations:** Department of Medical Microbiology, College of Health Sciences Makerere University; Department of Medical Biochemistry, College of Health Sciences Makerere University; Department of Immunology and Molecular biology, College of Health Sciences Makerere University; Division of Intramural Research, National Institute of Allergy and Infectious Diseases, National Institutes of Health, Bethesda, MD, USA; Uganda Cancer Institute

**Keywords:** Prostate cancer, Clinical characteristics, Primary management, Uganda Cancer Institute

## Abstract

**Background:** Prostate cancer is the second most common cancer among men in Uganda, with over 2086 incident cases in 2018. This study’s objective was to report the clinical characteristics and primary management of men diagnosed with prostate cancer at the Uganda Cancer Institute from 1^st^ January 2015 to 31^st^ December 2019.

**Methods:** Records from all men diagnosed with Prostate cancer at the Uganda Cancer Institute from 1^st^ January 2015 to 31^st^ December 2019 were reviewed. Clinical characteristics and primary treatment were recorded. Risk categorization was done using the European Society for Medical Oncology prostate cancer risk group classification.

**Results:** total of 874 medical records for men diagnosed with prostate cancer was retrieved. The median age was 70 years (interquartile range 64–77). In this study, 501 (57.32%) patients had localized disease. Among patients with localized disease, 2 (0.23%) were classified as low-risk, 5 (0.53%) as intermediate-risk, and 494 (56.52%) as high-risk. Three hundred seventy-three (373) patients had metastatic disease at diagnosis. Among patients with distant metastases, the most common site of metastases was bone 143 (16.36%), followed by spinal cord 54 (6.18%), abdomen 22 (2.52%), and lungs 14 (1.60%). Regarding the primary treatment options majority of the patients were on chemotherapy 384(43.94%) followed by hormonal therapy 336 (38.44%) and radiotherapy 127 (14.53%).

**Conclusion:** The majority of the patients diagnosed with prostate cancer at the Uganda Cancer Institute presented with advanced disease. The primary treatments were mostly chemotherapy, hormonal therapy, and radiotherapy. There is a need to improve prostate cancer screening in regional health care facilities and the communities to enhance early detection and management of prostate cancer.

## Introduction

Prostate cancer is the second most common cancer among men worldwide. In 2018, the Global Cancer Observatory (Globocan) report indicated 1,276,106 new prostate cancer cases, with approximately 358,989 deaths from the same disease (1). In Africa, prostate cancer is the most common cancer among men with a varying incidence in the different parts of the continent. The age-standardized prostate cancer incidence in eastern, western, northern, central, and southern parts of Africa is 23.9, 31.9, 13.2, 35.9, and 64.1 per 100,000 men, respectively (2,3). In Uganda, prostate cancer is the second most common cancer among men. In 2018, the incidence of prostate cancer in Uganda was 6.4%, with over 2086 new cases and 1177 deaths (1). Different studies have shown that African American men are at higher risk of prostate cancer than Caucasians counterparts (4,5). African American men with prostate cancer have also been found to have a worse prognosis and reduced survival; however, the underlying mechanisms are poorly understood (6). There is a close linkage in the genetics of prostate cancer in African American men and African men. African men with prostate cancer have been seen with aggressive disease and early presentation with an average age of 45-55 years (7). There is little research on prostate cancer in the African population, and this is further complicated by the limited number of functional cancer registries to capture accurate population data. We conducted a retrospective review of medical records of patients diagnosed with prostate cancer from 1st January 2015 to 31^st^ December 2019 at the Uganda Cancer Institute (UCI) to describe the clinical characteristics and primary management.

## Materials and Methods

Medical records of all men diagnosed with Prostate cancer at the UCI from 1^st^ January 2015 to 31^st^ December 2019 retrieved. The UCI is a national cancer center and is also designated as the East Africa center of excellence in cancer care, research, and treatment. The Uganda Cancer Institute, established in 1967, has a total bed capacity of 200 beds and daily patient attendance of 300. Person Identity Numbers were extracted from our local cancer register using the following words: Prostate cancer, Prostate adenocarcinoma, Prostate ductal carcinoma, or Prostate small-cell carcinoma. Subsequently, patient charts were reviewed. The following variables were registered: date of diagnosis, age, family history of prostate cancer, geographical region of origin, Gleason score, baseline PSA, TNM staging status, metastatic disease, organs involved, and treatment options. The presence of co-morbidities such as HIV, heart disease, diabetes, and hypertension, was recorded. In total, 1021 men were identified in the local cancer register search as diagnosed with Prostate cancer between 1st January 2015 and 31st December 2019 at the UCI. We extracted files and excluded 147 files that did not have a histological diagnostic report signed by a pathologist. Risk classification was performed according to the European Society for Medical Oncology (ESMO) guideline: Low risk: cT1-2a, PSA<10, GS_6; intermediate-risk: cT2b-c, PSA 10–20, GS 7; High-risk: T3-T4, PSA>20; Metastatic: N1, M1(8)(Table 3). The study was approved by the Makerere University, College of Health Sciences, School of Biomedical Sciences Research and Ethics Committee, and administrative clearance obtained from the UCI Research and Ethics Committee.

**Table 1:**
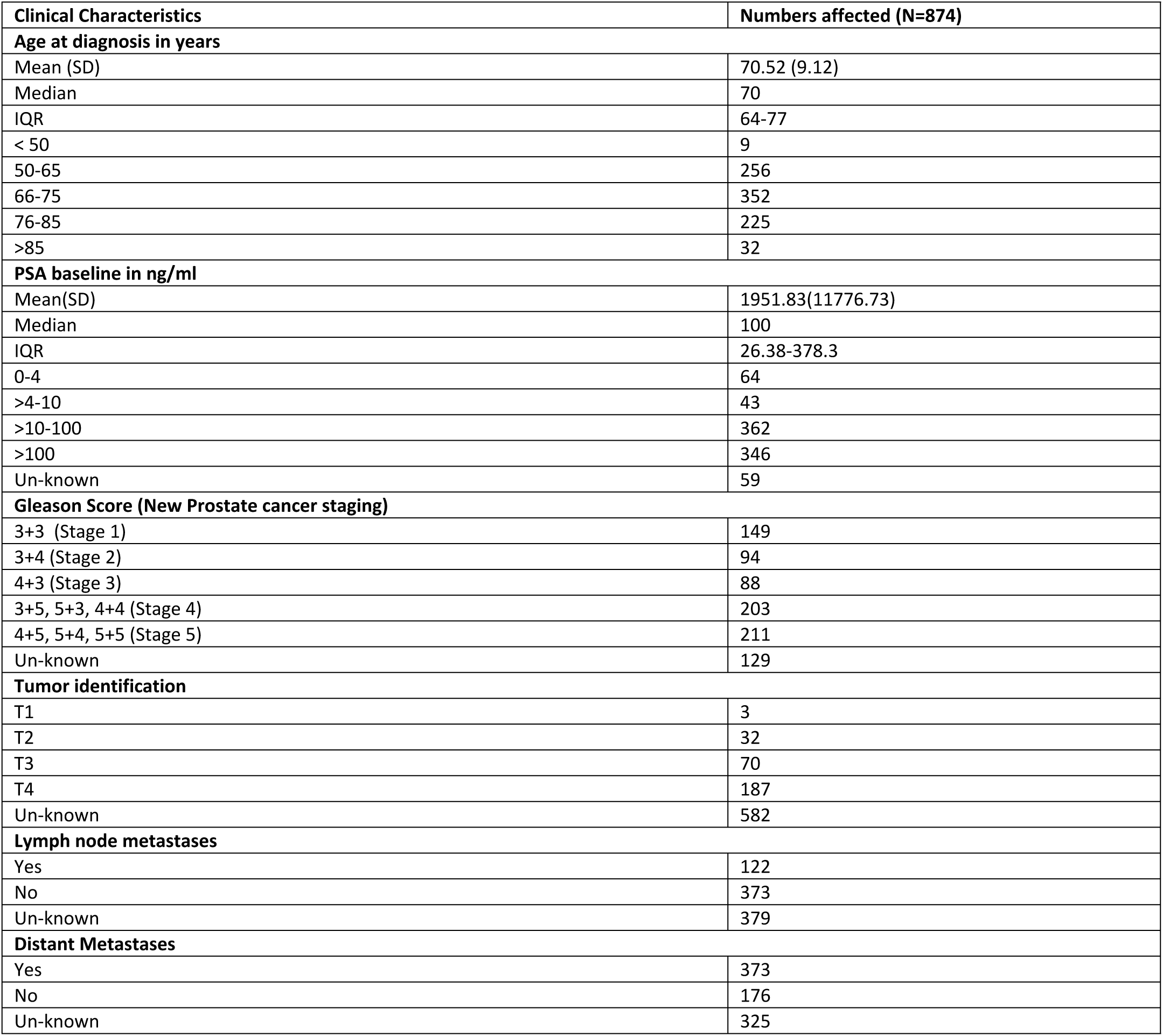
Clinical characteristics of Prostate cancer patients at the UCI from 2015 to 2019.

| Clinical Characteristics | Numbers affected (N=874) |
| --- | --- |
| **Age at diagnosis in years** |  |
| Mean (SD) | 70.52 (9.12) |
| Median | 70 |
| IQR | 64-77 |
| < 50 | 9 |
| 50-65 | 256 |
| 66-75 | 352 |
| 76-85 | 225 |
| >85 | 32 |
| **PSA baseline in ng/ml** |  |
| Mean(SD) | 1951.83(11776.73) |
| Median | 100 |
| IQR | 26.38-378.3 |
| 0-4 | 64 |
| >4-10 | 43 |
| >10-100 | 362 |
| >100 | 346 |
| Un-known | 59 |
| **Gleason Score (New Prostate cancer staging)** |  |
| 3+3 (Stage 1) | 149 |
| 3+4 (Stage 2) | 94 |
| 4+3 (Stage 3) | 88 |
| 3+5, 5+3, 4+4 (Stage 4) | 203 |
| 4+5, 5+4, 5+5 (Stage 5) | 211 |
| Un-known | 129 |
| **Tumor identification** |  |
| T1 | 3 |
| T2 | 32 |
| T3 | 70 |
| T4 | 187 |
| Un-known | 582 |
| **Lymph node metastases** |  |
| Yes | 122 |
| No | 373 |
| Un-known | 379 |
| **Distant Metastases** |  |
| Yes | 373 |
| No | 176 |
| Un-known | 325 |

**Table 2:**
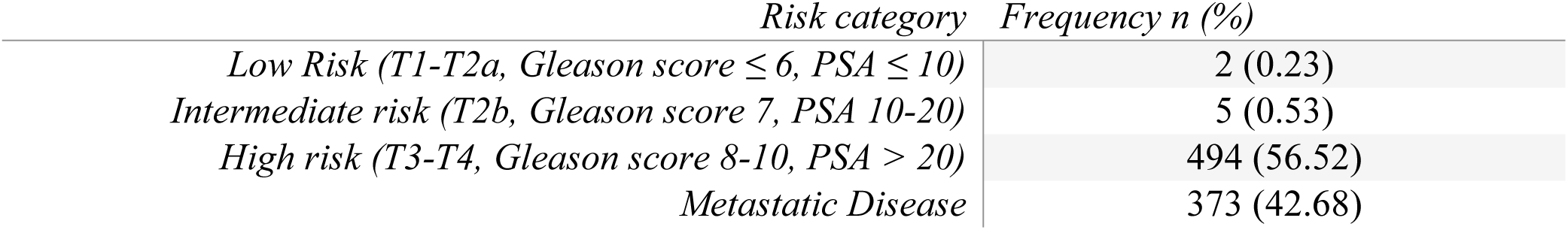
ESMO Prostate cancer risk categorization.

| *Risk category* | *Frequency n (%)* |
| --- | --- |
| *Low Risk (T1-T2a, Gleason score ≤ 6, PSA ≤ 10)* | 2 (0.23) |
| *Intermediate risk (T2b, Gleason score 7, PSA 10-20)* | 5 (0.53) |
| *High risk (T3-T4, Gleason score 8-10, PSA > 20)* | 494 (56.52) |
| *Metastatic Disease* | 373 (42.68) |

**Table 3:** Frequencies of patients on different treatment options.

| *Treatment options* | *Frequency n (%)* |
| --- | --- |
| *Surgery* | 78 (8.92) |
| *Chemotherapy* | 384 (43.94) |
| *Radiotherapy* | 127(14.53) |
| *Hormonal therapy* | 336(38.44) |
| *Palliative Care* | 83(9.50) |

## Results

We analyzed data of 874 men diagnosed with prostate cancer from 1^st^ January 2015 to 31^st^ December 2019 at the UCI. The majority of the patients were from the central part of Uganda 375 (42.12%), followed by western 237 (27.12%), eastern 182 (20.82%), and northern 67(7.67%), while 13 (1.49%) did not have documentation of area of residence. The median age was 70 years (interquartile range 64 to 77 years), the median baseline PSA was 100ng/ml (interquartile range 26.38 to 378.3 ng/ml), with over 39.6% (346) of the patients having a baseline PSA of >100ng/ml. Upon analysis of histological grading we observed the following Gleason scores; 6 in 17.05% (n=149), 7 in 20.82% (n=182), 8 in 23.23% (n=203) and 9-10 in 24.14% (n=211). In this group of patients, we noticed that 122 (13.96%) of the patients had lymph node metastases, and 373 (42.68%) had distant metastases. Among patients with distant metastases, the most common sites of metastases were bone 143 (16.36%), followed by spinal cord 54 (6.18%), abdomen 22(2.52%), and lungs 14(1.60%) among other sites. The summary of the clinical characteristics is, as shown in **Table 1**. A further analysis using the ESMO prostate cancer risk categorization, we found that 2 (0.23%), 5 (0.53%), 494 (56.52%), and 373 (42.68%) were having a low risk, intermediate risk, high risk, and metastatic disease respectively as shown in **Table 2**. Therefore, we observe that over 99.12% of prostate cancer patients admitted and managed at the UCI from 2015 to 2019 had advanced disease. Among patients with co-morbidities, 49 (5.6%) were HIV positive, 56 (6.4%) had diabetes mellitus, 154 (17.62%) had hypertension, and 33 (3.78%) had heart disease. Among the presenting complaints, the most common presenting complaint was lower urinary tract symptoms 306 (35.01%), followed by bone pain 60 (6.86%), among other symptoms. Regarding the treatment options provided to these patients at the UCI, 384 (43.94%), 336 (38.44%), 127 (14.53%), 78 (8.92%) and 83 (9.50%) patients received chemotherapy, hormonal therapy, radiotherapy, surgery, and palliative care respectively as shown in **Table 3**. The most commonly used chemotherapeutic agents are androgen deprivation agents, mainly bicalutamide and a taxane, docetaxel. Most patients received diethylstilbestrol (DES), a synthetic ethinyl estrogen as the main choice for hormonal therapy.

## Discussion

In sub-Saharan Africa, prostate cancer morbidity and mortality are high, attributed to weak national cancer screening structures and the late presentation of patients (9). The Globocan report of 2018 indicates that Uganda had 2086 (6.4%) incident cases of prostate cancer, and this was the second most common cancer in males (1). In Uganda, more than 90% of all the prostate cancer cases seen at the UCI are referrals from the region, and these often present with advanced disease (10). Few studies in sub-Saharan Africa have described the clinical and demographic patterns of prostate cancer patients. In this study, we described the clinical presentation, demographics, and treatment given to prostate cancer patients at the UCI in Kampala Uganda from 2015 to 2019.

In this study, most patients were managed on chemotherapy, hormonal therapy, and radiotherapy. This is because most of the patients present with advanced disease, and hence surgery and watchful waiting were un-common treatment options. The access to cancer treatment services is a big challenge in Uganda since most patients have to travel long distances to the UCI the main oncology treatment facility. This is compounded by the limited human resource and the lack of technologically advanced treatment modalities like radiotherapy machines and immunotherapy. In Uganda, there is one cobalt 60 machine currently serving all the cancer patients in a country of over 42 million people.

There is a scarcity of second and third generation androgen deprivation chemotherapeutic agents such as enzalutamide that are more efficacious in the management of advanced prostate cancer in the country. This is also due to the high cost of these agents. In this study, most of the patients on chemotherapy received bicalutamide, the agent provided on the government drug list. There were a small number of patients that underwent surgery (bilateral orchiectomy). This could be explained by the limited number of urologists in the country that can carry out such a surgical procedure or the number of patients willing to undergo the procedure. Several studies indicate that prostate cancer disease pathogenesis is associated with genetic and environmental factors. Studies in African American men have shown that particular genetic polymorphisms are associated with reduced prostate cancer disease outcomes. Studies done in African men with prostate cancer indicate a close association with the same gene loci seen to be associated with prostate cancer risk in African American men (11,12). These findings could explain the advanced disease presentation for more than 99% of men with prostate cancer. The major limitation of this study was missing information due to poor record-keeping, and this could have affected analysis. Therefore, we recommend improving prostate cancer screening programs in the country with an emphasis on early cancer detection. There is also a need to decentralize the screening, diagnosis, and treatment of cancers in the country through the establishment of regional treatment centers. There is a need to invest in advanced treatment facilities like next-generation radiotherapy machines, the introduction of immunotherapy, and more research into understanding the etiology and pathogenesis of prostate cancer in the region.

## Conclusion

In this study, we describe clinical and demographic characteristics among Ugandan men with prostate cancer and understand the risk stratification and primary treatment. The majority of the men were diagnosed with advanced disease and primarily managed with chemotherapy, hormonal therapy, and radiotherapy.

## Authors’ contributions

PK, OJS, MJ, SJR, and NN contributed to the conceptualization of the manuscript. MY contributed to data abstraction and analysis. PK drafted the collected, cleaned, analyzed data, and drafted the manuscript. All authors reviewed the manuscript for publication. All authors read and approved the final manuscript.

## Acknowledgments

None to declare

## Availability of data and materials

All data files used to in this manuscript article [and its supplementary information files] are available via figshare DOI; <u>10.6084/m9.figshare.12610727</u>

## Consent for publication

Not applicable.

## Ethics approval and consent to participate

The study was approved by the Makerere University, College of Health Sciences, School of Biomedical Sciences Research and Ethics Committee, and administrative clearance was obtained from the UCI Research and Ethics Committee.

